# Dual-located WHIRLY1 affects salicylic acid homeostasis *via* coordination of ICS1, PAL1 and BSMT1 during *Arabidopsis* plant aging

**DOI:** 10.1101/2020.05.04.077388

**Authors:** Wenfang Lin, Hong Zhang, Dongmei Huang, Dirk Schenke, Daguang Cai, Binghua Wu, Ying Miao

**Author notes:** Corresponding author Fujian Provincial Key Laboratory of Plant Functional Biology, Fujian Agriculture and Forestry University, Fuzhou 350002, China. The author responsible for distribution of materials integral to the findings presented in this article in accordance with the policy described in the Instructions for Authors is: Ying Miao, Binghua Wu. Y.M. designed the study. W.F.L. performed SA measurements, western blots, phenotyping, and qRT-PCR. D.H. performed, ChIP-seq, ChIP-qPCR, H.Z. performed plasmid constructs and promoter activation activity and the mutants screening. B.H.W performed microarray data analyses. W.F.L. and Y.M. analyzed the data. Y.M. and D.S. wrote the paper. No conflict of interest.

## Abstract

Salicylic acid (SA) homeostasis determines also developmental senescence and is spatiotemporally controlled by various mechanisms, including biosynthesis, transport and conjugate formation. The alteration of WHIRLY1 (WHY1), a repressor of leaf natural senescence, with respect to allocation in the nucleus or chloroplast causes a perturbation in SA homeostasis, resulting in adverse plant senescence phenotypes. Loss of *WHY1* resulted in a 5 days earlier SA peak compared to wild type plants which accumulated SA at 42 days after germination. SA accumulation coincided with an early leaf senescence phenotype, which could be prevented by ectopic expression of the nuclear WHY1 isoform (nWHY1). However, expressing the plastid WHY1 isoform (pWHY1) greatly enhanced cellular SA levels. A global transcriptional analysis in WHY1 loss-of-function background by expressing either pWHY1 or nWHY1 indicated that hormone metabolism related genes were most significantly altered. The pWHY1 isoform predominantly affected stress related gene expression, while the nWHY1 controlled rather developmental gene expression. Chromatin immunoprecipitation-qPCR (ChIP-qPCR) assays indicated that nWHY1 directly binds to the promoter region of isochorismate synthase (*ICS1)* to activate *its* expression at later stage, but indirectly activated S-adenosyl-L-methionine-dependent methyltransferase (*BSMT1)* gene expression *via* ethylene response factor 109 (ERF109), while repressing phenylalanine ammonia lyase (*PAL1)* expression *via* R2R3-MYB member 15 (MYB15) at the early stage of development. Interestingly, rising SA levels exerted a feedback effect by inducing nWHY1 modification and pWHY1 accumulation. Thus, the alteration of WHY1 organelle isoforms and the feedback of SA intervened in a circularly integrated regulatory network during developmental or stress-induced senescence in *Arabidopsis*.

## Introduction

Salicylic acid is crucial for plant growth, responses to pathogens, e.g. by programmed cell death and environmental responses. Its homeostasis is temporally and spatially controlled by various mechanisms, including biosynthesis, transport and conjugate formation. For example, leaf development in *Arabidopsis* was regulated by SA biosynthetic / signaling genes. Early leaf senescence is a result of SA overproduction in mutants such as isochorismate synthase (ICS1) and phenylalanine ammonia lyase (PAL) overexpression lines (Love et al., 2008; Rivas-San et al., 2011), whereas the hypersensitive response (a fast form of programmed cell death) have been intensively investigated in the S-adenosyl-L-methionine-dependent methyltransferase (*bsmt1)* mutant (Vlot et al., 2009). There are two main SA biosynthetic pathways in plants: the phenylalanine ammonia lyase (PAL) pathway and the isochorismate (IC) pathway, both depending on the primary metabolite chorismate (Dempsey et al. 2011). In the PAL pathway, the chorismate-derived L-phenylalanine is converted into SA *via* either benzoate intermediates or coumaric acid through a series of enzymatic reactions involving PAL, benzoic acid 2-hydroxylase (BA2H), and other uncharacterized enzymes (Leon et al. 1995b). Approximately 10% of defense-related SA is produced by the cytosolic PAL pathway and in Arabidopsis four PAL enzymes have been identified. In the IC pathway, chorismate is converted in a two-step process to SA *via* isochorismate involving isochorismate synthase (ICS) and isochorismate pyruvate lyase (IPL). In *Arabidopsis*, two ICS enzymes have been described to convert chorismate to isochorismate, but in recent studies another isochorismate synthase was identified (Rekhter et al. 2019; Torrens-Spence et al. 2019). This pathway accounts for ∼90% of the SA production generated by the plastid-localized ICS1 inducible by pathogens and UV light (Wildermuth et al. 2001; Garcion et al. 2008). Endogenous SA undergoes a series of chemical modifications including hydroxylation, glycosylation, methylation and amino acid conjugation. These modifications directly affect the biochemical properties of the SA derivatives, and play a pivotal role in SA catabolism and homeostasis to regulate leaf senescence (Zhang et al. 2013). It has been shown that SA affects regulation of gene expression during leaf senescence (Morris et al. 2003; Vogelmann et al.2013; Zhang et al. 2013; 2017) and in advancing flowering time in *Arabidopsis thaliana* (Martínez et al. 2004), as well as in inhibiting seed germination (Alonso-Ramirez et al. 2009; Lee et al. 2013). Although SA biosynthesis and its function in both local and systemic acquired resistance (SAR) against microbial pathogens and in plant development were well understood (Park et al. 2007; An and Mou, 2011), the underlying molecular mechanism of free SA homeostasis in cells is less clear.

WHIRLY family proteins are dually located in both the nucleus and organelles, and perform numerous cellular functions in both locations (Krause et al. 2005; Grabowski et al. 2008). In the nucleus, WHIRLY1 (WHY1) protein was found to regulate the expression of genes related to defense and senescence by binding to their respective promoters (Desveaux et al. 2000; Desveaux et al. 2004; Xiong et al. 2009; Miao et al. 2013; Krupinska et al. 2013). WHY1 protein binds for example to the promoter of *WRKY53* and repress *WRKY53* and *WRKY33* expression in a development-dependent manner during early senescence in *Arabidopsis* (Miao et al. 2013; Ren et al. 2017), while it activates the *HvS40* gene during natural and stress-related senescence in barley (*Hordeum vulgare*) (Krupinska et al. 2013) and *PsbA* gene expression in response to chilling treatment in tomato (Zhuang et al. 2018). In the nucleus, WHY1 protein also modulates telomere length by binding to their *AT*-rich region (Yoo et al. 2007) and affects microRNA synthesis (Swida-Barteczka et al. 2018). Moreover, in chloroplasts, WHY1 has a function on organelle genome stability, facilitating accurate DNA repair (Cappadocia et al. 2010, 2012; Lepage et al. 2013) and affects RNA editing/splicing (Prikryl et al., 2008; Melonek et al. 2010). The intracellular localization of WHY1 and/or the developmental stage of the plants may contribute to its various functions (Ren et al. 2017). Furthermore, WHY1 has been reported to be involved in (a)biotic stress signaling pathways, e.g. in response to chilling (Zhuang et al. 2018), high light (Kucharewicz et al. 2017), N deficiency (Comadira et al. 2013), reactive oxygen species (Lin et al. 2019; Lepage et al. 2013), hormones such as SA and abscisic acid (Xiong et al. 2009; Isemer et al. 2012) and defense signaling, being e.g. required for SA- and pathogen-induced PR1 expression (Desveaux et al. 2005).

In this study, we extend the roles of the dual-located WHY1 protein with respect to SA biosynthesis *via* regulating *PAL1* and *ICS1* expression and SA modification *via* affecting *BSMT1* gene expression, in a developmental dependent manner. Moreover, the cellular SA level affected the distribution and status of WHY1 protein in the nucleus and in plastids, suggesting a feedback mechanism to regulate SA homeostasis. Further, globally analysis of gene expression in loss-of WHY1 and gain-of pWHY1 or nWHY1 indicated that the levels of hormone metabolism related genes were significantly altered. Our results provide the first evidence that the dual-located WHY1 protein exerts a novel function in both nucleus and chloroplasts to fine-tune SA homeostasis affecting plant aging in *Arabidopsis*.

## Results

### WHY1 changes the gene expression level of *PAL, ICS* and *BSMT1* and SA contents during plant aging

To explore how WHY1 involves in the SA metabolism pathways (Figure 1a), we used the *why1-1* mutant previously deployed in several of our studies (Miao et al. 2013; Ren et al. 2017; Lin et al. 2019). This *why1-1* mutant displays an early senescence phenotype (Miao et al. 2013), similar to the S-adenosyl-L-methionine-dependent methyltransferase (*bsmt1*) mutant (Vlot et al. 2009) and the SA 3-hydroxylase (*s3h*) mutant (Zhang et al. 2013). We analyzed the expression levels of *ICS, PAL, BSMT1*, encoding a protein with both benzoic acid (BA) and SA carboxyl methyltransferase activities, and salicylic acid glucoside/glucose ester modification enzymes such as *UGT71B1, UGT89B1* or *UGT74F2* (Dempsey et al. 2011) in the *why1* mutant compared to WT during plant development from 28 to 42 days after germination (dag). Interestingly, loss-of-*WHY1* increased the transcript level of *PAL1* and *PAL2* at 37 dag, but greatly decreased the transcript level of *BSMT1* at 35 and 37 dag and of *ICS1* at 42 dag (Figure 1b), while the transcript levels of *UGT71B1, UGT74F2, UGT89B1* and *S3H* were not altered in the *why1* mutant during plant development (Supplementary Fig S1).

**Figure 1.**
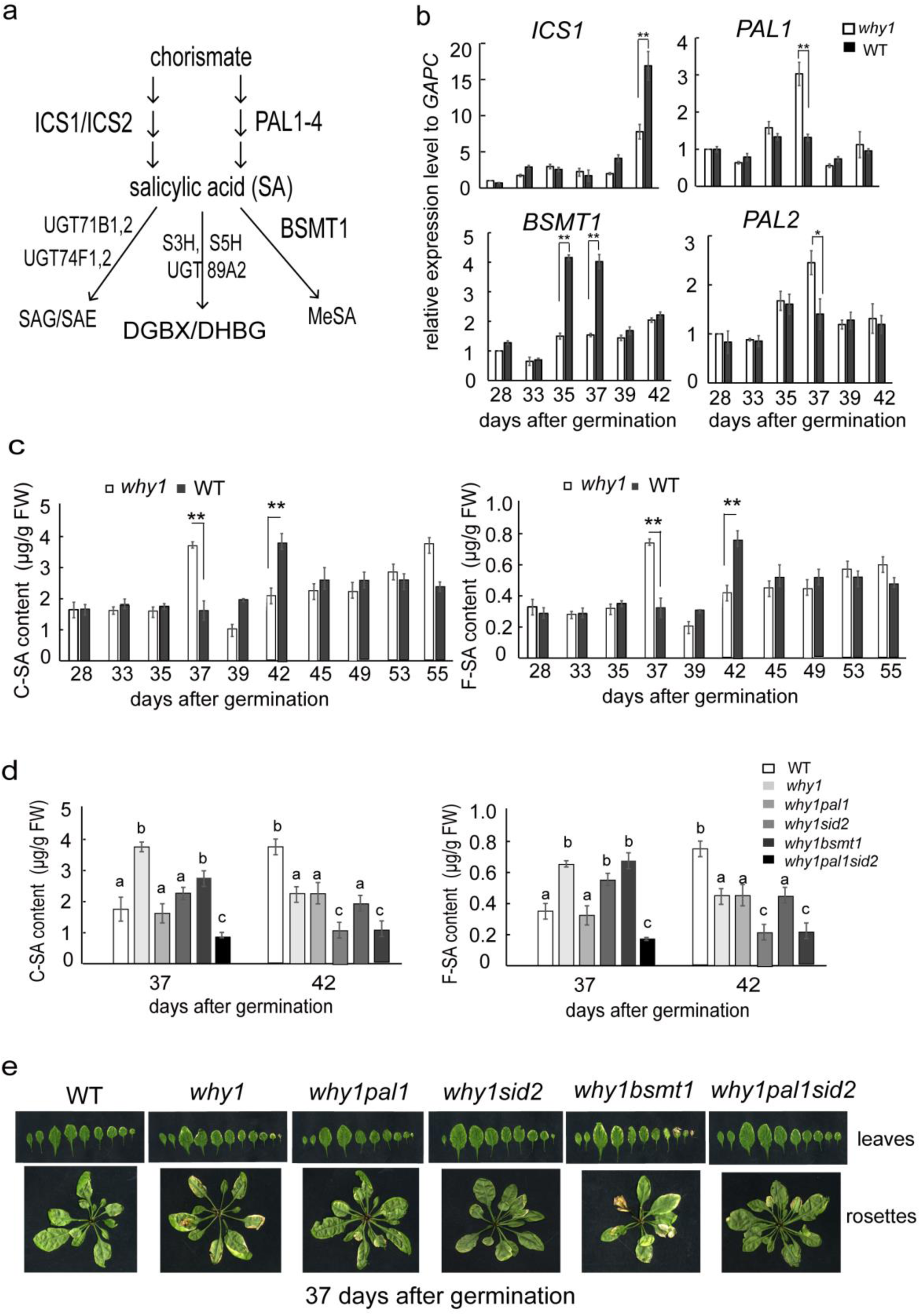
The variation transcript level of genes encoding key enzymes related to SA metabolism pathway and SA contents in the *why1* line during the development a. SA metabolism pathway in the cell. b The variation transcript level of genes encoding key enzymes related to SA metabolism in the *why1* line during plant development. c. Content of conjugated (C-SA) and free (F-SA) salicylic acid in wild type and *why1* mutant during the period of 28 to 55 days after germination (dag); d. Changes of conjugated and free salicylic acid contents in a series of double mutants with focus on 37 and 42 dag. f. Senescence phenotype of 37 dag old double mutants. The relative expression level normalized to *GAPC*, wild type at 28 dag (b) was setup as 1. The standard error bars present three time biological replicates and three time techniques replicates, the values are shown as means ±SD. Asterisks (*P < 0.05, **P < 0.01) show significant differences to wild type line according to either two-way ANOVA or pair-wide multiple t-tests.

Thus, we tested whether SA contents also changed in the *why1* mutant during plant aging. The SA contents including conjugated and free type of SA of the *why1* and WT plants were measured with a HPLC assay during the period from 28 dag to 58 dag of plant development. Our results indicate that *loss-of-WHY1* made both conjugated SA and free SA peak 5 days earlier (at 37 dag) than in wild type (Figure 1c-d).

In order to genetically confirm this hypothesis, we produced t*he why1pal1, why1sid2, why1pal1sid2, why1bsmt1* double/triple mutants (Supplementary Fig S2) and measured the SA contents in these mutants during plant aging (Figure 1e). Interestingly, the early SA peak disappeared in the *why1pal1* line at 37dag, showing a similar SA profile as the wild type, while SA accumulation in *why1* mutants combined with *bsmt1* mutation were not that strongly affected, displaying the same early senescent phenotype as the *why1* line. However, SA accumulation in *why1* combined with *sid2* (ics1) was inhibited at 42 dag. The *whylpal1sid2* triple mutant showed a delay senescence phenotype and had again no earlier SA peak even maintain low level of SA at 37 and 42 dag during plant development, suggesting that PAL activity is crucially important for SA accumulation at early stage. Thus, we genetically confirmed that SA homeostasis in cells is affected by WHY1 predominantly by its effect on PAL1.

### nWHY1/pWHY1 affects the gene expression level of *PAL1, ICS1* and *BMST1* as well as SA homeostasis during plant aging

As we knew, WHY1 is dual-located in the nucleus and plastids (Grabowski et al. 2008). To clarify which isoform of WHY1 affects SA metabolism and its homeostasis, we complemented the *why1* background line with pWHY1,nWHY1 and pnWHY1 under *35S* promoter control (Lin et al. 2019), and analyzed the transcript levels of *PAL, ICS, and BSMT1* from 28 dag to 42 dag. Complementation with nWHY1 or full length WHY1 (pnWHY1) restored wild type transcript levels of *PAL1, PAL2* and *BSMT1*, while the *nWHY1/why1* line had even lower *PAL1* expression level at 37 dag and 42 dag compared to WT. Surprisingly, complementation with pWHY1 not only pronounced the transcript level of *PAL1* two folds and repressed the transcript level of *BSMT1* at 37 dag, but also significantly increased the transcript level of *ICS1* at 42 dag, (Figure 2a). Measuring the SA contents in the complemented *why1* mutant background from 28 to 42 dag, both *nWHY1/why1* and *pnWHY1/why1* lines significantly restored wild type SA accumulation of the *why1* line until 37 dag and in the *nWHY1/why1* mutant the SA content was even lower at 42 dag. However, pWHY1 significantly pronounced SA accumulation during the whole period of development (Figure 2b), indicating that nWHY1 somehow repressed SA accumulation via suppression of *PAL1* expression. On the other hand, pWHY1 might pronounce SA accumulation via repressing *BSMT1* during the early stages and promoting *ICS1* at the late stage, in a developmentally dependent manner.

**Figure 2.**
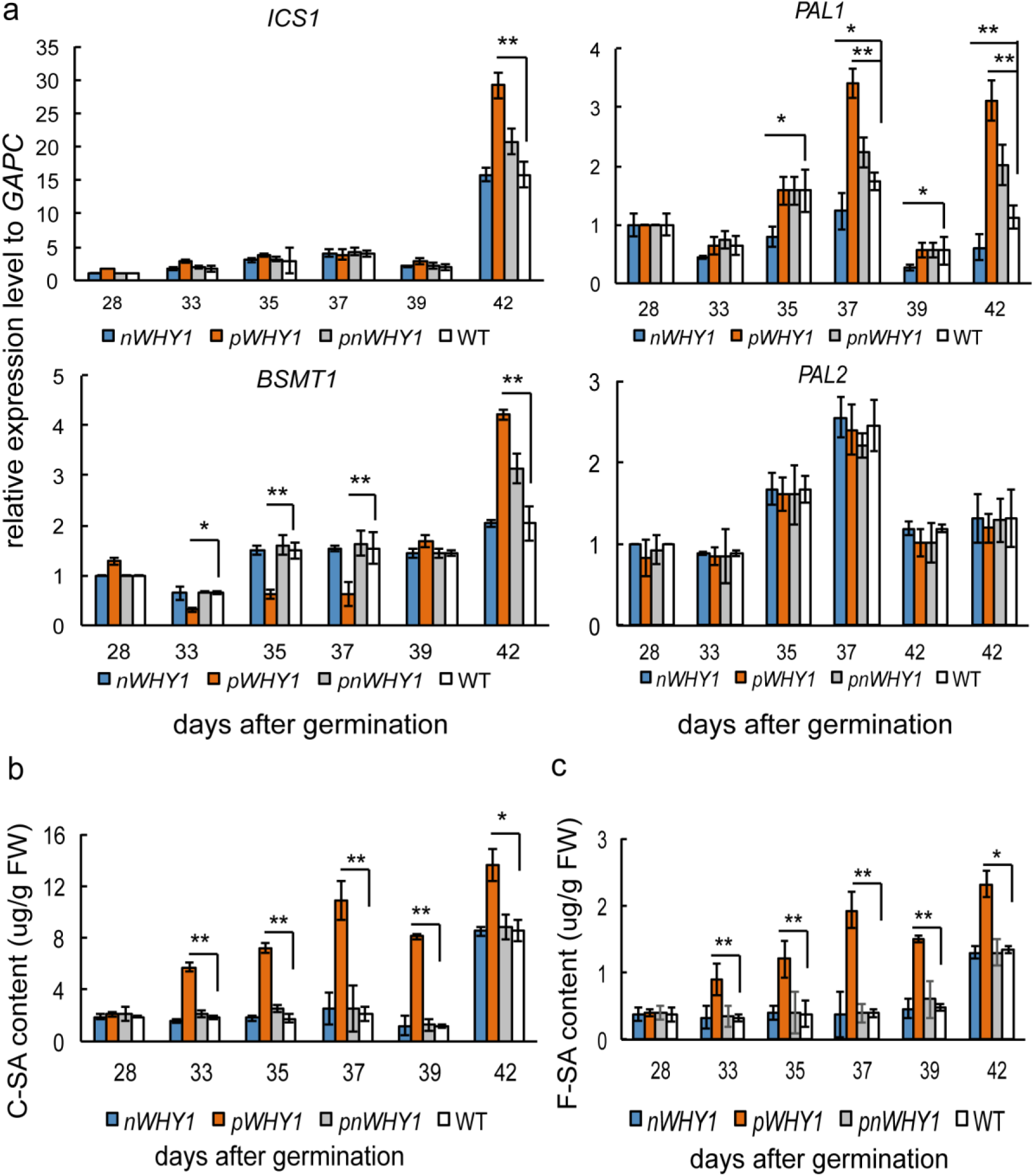
Transcript level analysis of genes encoding key enzymes related to SA metabolism pathway (a) and SA contents (b) in the *pWHY1/why1, nWHY1/why1*, and *pnWHY1/why1* transgenic plants compared to wild type from 28 to 42 dag during plant development.

The standard error bars present three time biological replicates, the values are shown as means±SE. Asterisks (*P < 0.05, **P < 0.01) show significant differences to WT within the respective conditions according to Student’s t test.

### Hormone- related gene enrichment in “compartmental WHY1” transgenic plants

In order to globally understand the differences and similarities of the nuclear transcriptome response between pWHY1 and nWHY1, a microarray sequencing analysis was deployed. Phenotypic differences were observed in the short term response and to avoid a long term secondary artifact caused by continuous expression, an estradiol-inducible promoter was used to generate “inducible compartmental WHY1” transgenic plants (*VEX:pWHY1/why1* and *VEX:nWHY1/why1*) as described in Ren et al., (2017). We found that WHY1 protein level increased about 14 folds after two hours induction with 20 μM estradiol (Ren et al. 2017). The total RNA isolated from the 5 week old rosette leaves of inducible *VEX:pWHY1/why1* and *VEX:nWHY1/why1* plants before (0h) and after estradiol application (2h), as well as from *why1* and WT plants was used for transcriptome analysis by ATH1 *Arabidopsis* GeneChip microarrays with two biological replicates. Comparing the transcriptome of inducible pWHY1 plants to that of non-inducible pWHY1 plants revealed a complex genetic reprogramming with 1165 and 4560 transcripts being at least 2-fold up- and down-regulated, respectively. Comparison of inducible nWHY1 plants to that of non-inducible nWHY1 plants revealed also a complex genetic reprogramming with 920 and 3965 transcripts up- and down-regulated, respectively. Transcriptomic comparison of the *why1* mutant to WT plants identified 4432 and 1190 transcripts up- and down-regulated, respectively (Supplementary Fig S3).

To visualize gene expression reprogramming in the *VEX:pWHY1 VEX:nWHY1* and the *why1* plants, their entire nuclear transcriptome was subjected to MapMan analysis allowing the identification of biological processes with significant alterations (Thimm et al., 2004). The hormone metabolism pathways are significantly overrepresented after induction of pWHY1, nWHY1, or by loss-of WHY1, affecting especially auxin, jasmonic acid (JA) and ethylene metabolism, as well as SA metabolism (Figure 3, Supplemental dataset1-4). The regulation of secondary metabolism and stress are also significantly enriched after induction of pWHY1 expression (Figure 3a). These stresses are associated with biotic stresses and abiotic stresses responses that are related to redox imbalance. They mostly are up-regulated by pWHY1 (Figure 3a). In contrast the regulation of RNA, development and signaling terms are significantly enriched after induction of nWHY1 expression. Since the opposite regulation of signaling, development, RNA and transport terms is observed in loss-of WHY1 plants (Figure 3a), these changes can be attributed to the inducible expression of pWHY1 or nWHY1 (Figure 3a). Globally, a net enrichment of biological processes linked to hormone metabolism is found within the most significantly differential expressed genes after induction of pWHY1 or nWHY1 or deletion of WHY1 (Figure 3b); a net enrichment for biological processes linked to hormone metabolism, secondary metabolism and photosynthetic stress is found within the most differentially expressed genes in inducible pWHY1 line (Figure 3b), while a net enrichment for biological processes linked to RNA regulation, development or signaling is found within the most differentially expressed genes in inducible nWHY1 line (Figure 3b) and a net enrichment for biological processes linked to photosynthesis and signaling or development or RNA regulation is found within the most differentially expressed genes in the *why1* line (Figure 3b).

**Figure 3.**
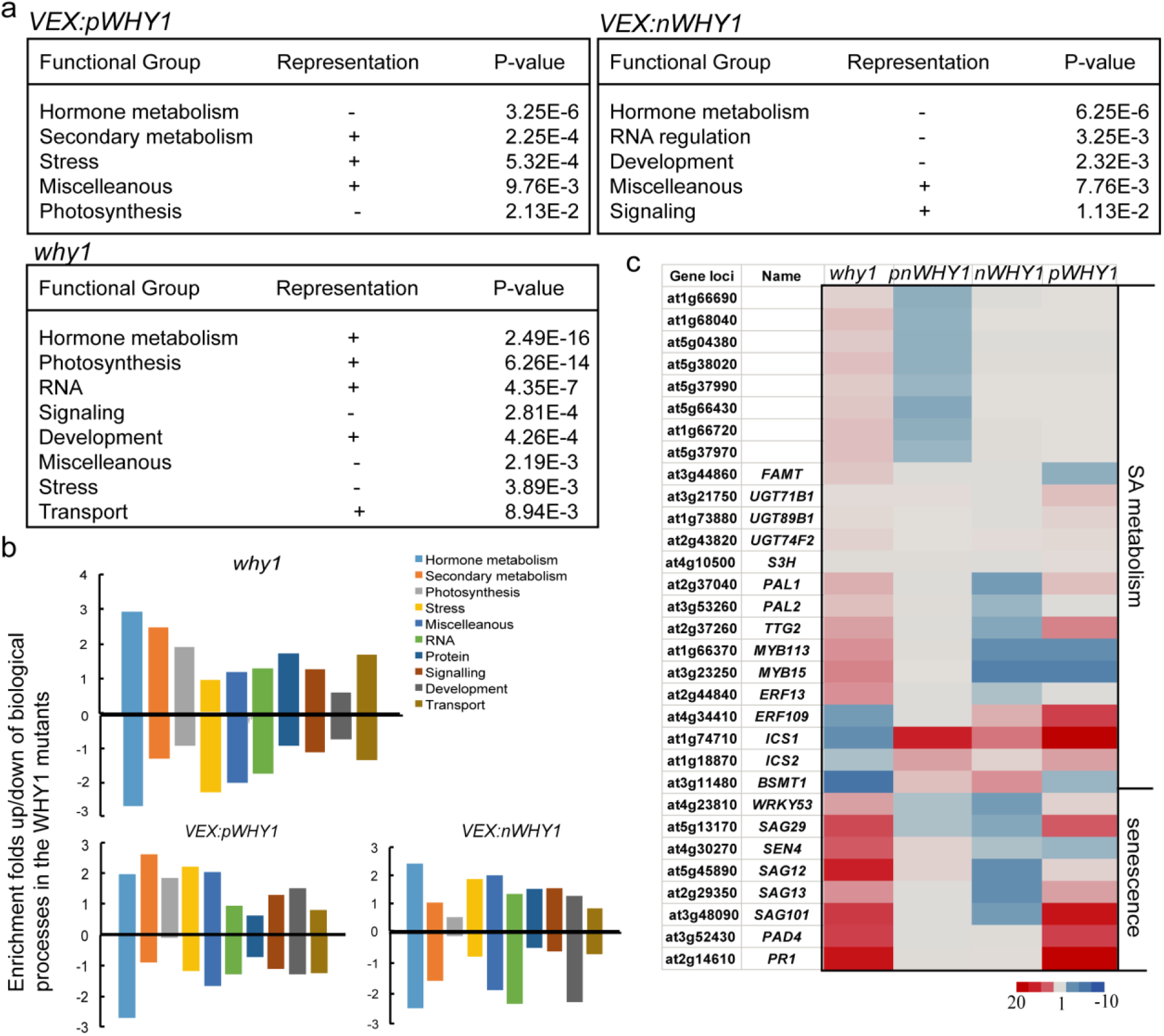
The V*EX:pWHY1, VEX:nWHY1* and the *why1* mutants exhibits a complex nuclear genetic reprogramming. a. MapMan analysis for gene ontology terms enrichment of the entire *VEX:pWHY1, VEX:nWHY1* and the *why1* nuclear transcriptome. b. Histogram presenting the ratio of differentially expressed genes enrichment changes of selected biological process of the *VEX:pWHY1, VEX:nWHY1* and the *why1* transcriptome. c. The heatmap of SA metabolism related gene expression levels of the *pWHY1/why1, nWHY1/why1, pnWHY1/why1* plants, and the *why1* mutants. *VEX:pWHY1, VEX:pWHY1/why1; VEX:nWHY1, VEX:nWHY1/why1*

Among the differentially expression genes, 153 of differentially expression genes overlay between inducible pWHY1 and nWHY1 lines. Among them, 42 of hormone-related gene expressions were up- or down-expression in the pWHY1 or *why1* lines, including SA, JA, IAA and ethylene metabolism and signaling related genes (Figure 3, Supplementary dataset1-4). The 24 highest expressed or suppressed genes in the pWHY1, nWHY1 or the *why1* plants, which encode key components of the SA metabolism pathway including ICS1, ICS2, PAL1, PAL2, UGT71B1, UGT89B1, UGT74F2, BSMT1, as well as SA signaling related genes, or senescence / cell death related genes are shown in the heatmap (Figure 3c).

### WHY1 directly binds at the promoter region of *ICS1* and indirectly affects *PAL1* and *BSMT1* expression in a developmental dependent manner

WHY1 was first reported as a transcription factor in the nucleus (Marechal et al. 2000). To investigate whether WHY1 directly regulates *ICS1, PAL1/PAL2, BSMT1* gene expression, we analyzed our previous ChIP-seq dataset and above microarray dataset, and found that *ICS1, MYB15* and *ERF109* are direct targets of WHY1 (Miao et al. 2013; and Figure 4a), but *PAL1* and *BSMT1* are not. A search for transcription factor binding motifs in promoter regions of *ICS1, MYB15, ERF109, PAL1*, and *BSMT1* genes was conducted with PlantCARE (Lescot et al. 2002) and resulted in two w-boxes, six MYC elements, and four MYB motives in the promoter of *PAL1;* 6xERE elements in the *BSMT1* promoter (Figure 4b) as well as several GTNNNNAAT and AT-rich motives in the *ICS1, MYB15*, and *ERF109* promoters. In order to clarify the relationship among them, firstly, we confirmed WHY1 binding at the target genes by chromatin immunoprecipitation qPCR (ChIP-qPCR) using leaf material from 37 and 42 dag of expressing HA-tagged WHY1 under its native promoter (*P*_*why1*_:*WHY1-HA)* as described in previous work (Miao et al. 2013). The putative cis elements found in *WRKY53, ICS1, MYB15, ERF109*, and *WRKY33* promoters, included several GTNNNNAAT or AT-rich motives (Figure 4b), and were enriched 5-20 fold (Figure 4c). The regions containing GTNNNNAAT and AT-rich motives of *MYB15, ERF109*, and *WRKY53* were enriched 10-15 folds at 37 dag, while fragments of *ICS1* and *WRKY33* could not be detected at 37 dag, but together with *MYB15* and *WRKY53* a high enrichment was observed at 42 dag (Figure 4c). Furthermore, the expression levels of these genes were analyzed by quantitative reverse transcription PCR (qRT-PCR) at 37 and 42 dag in *why1* and WT plants. WHY1 binding negatively correlated with gene expression in the knockout background of *ERF109* at 37 dag and ICS1 at 42 dag and positively with *MYB15* expression at both 37 and 42 dag. While *WRKY53* expression is positively correlated in *why1* plants at 37 dag, *WRKY33* was up-regulated at 42 dag. Thus, WHY1 appears to exert either negative effects on gene expression (WRKY53, WRKY33 and MYB15) or causes activation of its target genes, such as *ERF109* and *ICS1* depending on the developmental stage.

**Figure 4.**
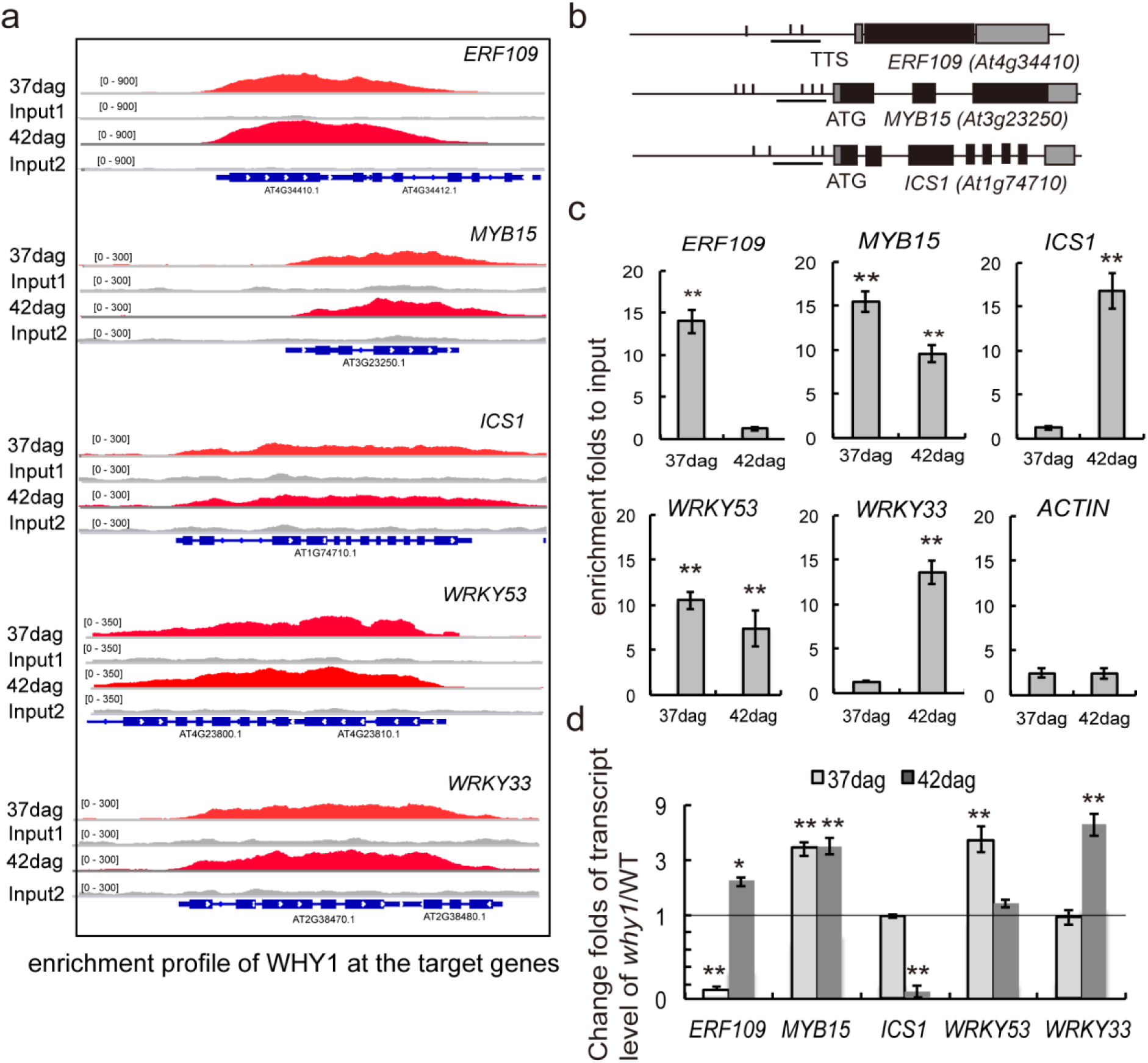
WHY1 activates/represses target gene expression a. Enrichment profiles of WHY1 protein in five target genes: *ERF109, MYB15, WRKY33, ICS1*, and *WRKY53* by ChIP-seq; b.Position of promoter motives (*GTNNNNAAT plus AT-rich*) of WHY1 target genes; c. Enrichment folds of WHY1 at the promoters of target genes by ChIP-qPCR at 37 and 42 days after germination; d. The expression levels of target genes at 37 and 42 days after germination in the *why1* mutant compared to WT. The error bars represented SD from three biological replicates. Asterisks indicated significant differences from the *ACTIN* according to two-tail Student’s t test (* denotes P < 0.05, ** for P < 0.01).

In order to further verify the activation or repression activity of WHY1, the promoter sequences of *WRKY53, ICS1, MYB15, ERF109, PAL1* and *BSMT1* were cloned into dual-luciferase vectors and applied in a transient expression assay using *Nicotiana benthamiana* leaves (Hellens et al., 2005). In addition to measure promoter activation or repression by WHY1, also MYB15, and ERF109 were included in the analysis to investigate indirect effects of WHY1 in the nucleus. The coding sequences of WHY1, MYB15 and ERF109 were cloned under the control of the Arabidopsis *ACTIN1* promoter (*ACTIN:WHY1-HA, ACTIN:MYB15-HA*, and *ACTIN:ERF109-HA*) (Figure 5a), and co-infiltrated with the reporter vector to drive LUCIFERASE (LUC) expression (Hellens et al., 2005). We then measured the LUC and RENNILASE (REN) luminescence ratio (i.e. LUC/REN ratio) in infiltrated leaves. To assess any basal activation or repression of putative promoters, a mini-*GAL4* promoter vector was used in each co-infiltration experiment as a control; the *WRKY53* promoter was used as a positive control. The results showed that WHY1 activated promoters of *ICS1* and *ERF109*, but it repressed the promoters of *MYB15* and *WRKY53* displaying the opposite expression pattern of the why1 knockout plants (Figure 4b). The transcription factors MYB15 and ERF109 were able to activate *PAL1, PAL2* and *BSMT1* gene expression, respectively (Figure 5b-c). Therefore, WHY1 directly activated *ICS1* expression and indirectly affected *PAL1, PAL2* and *BSMT1* gene expression *via* MYB15 and ERF109, respectively.

**Figure 5.**
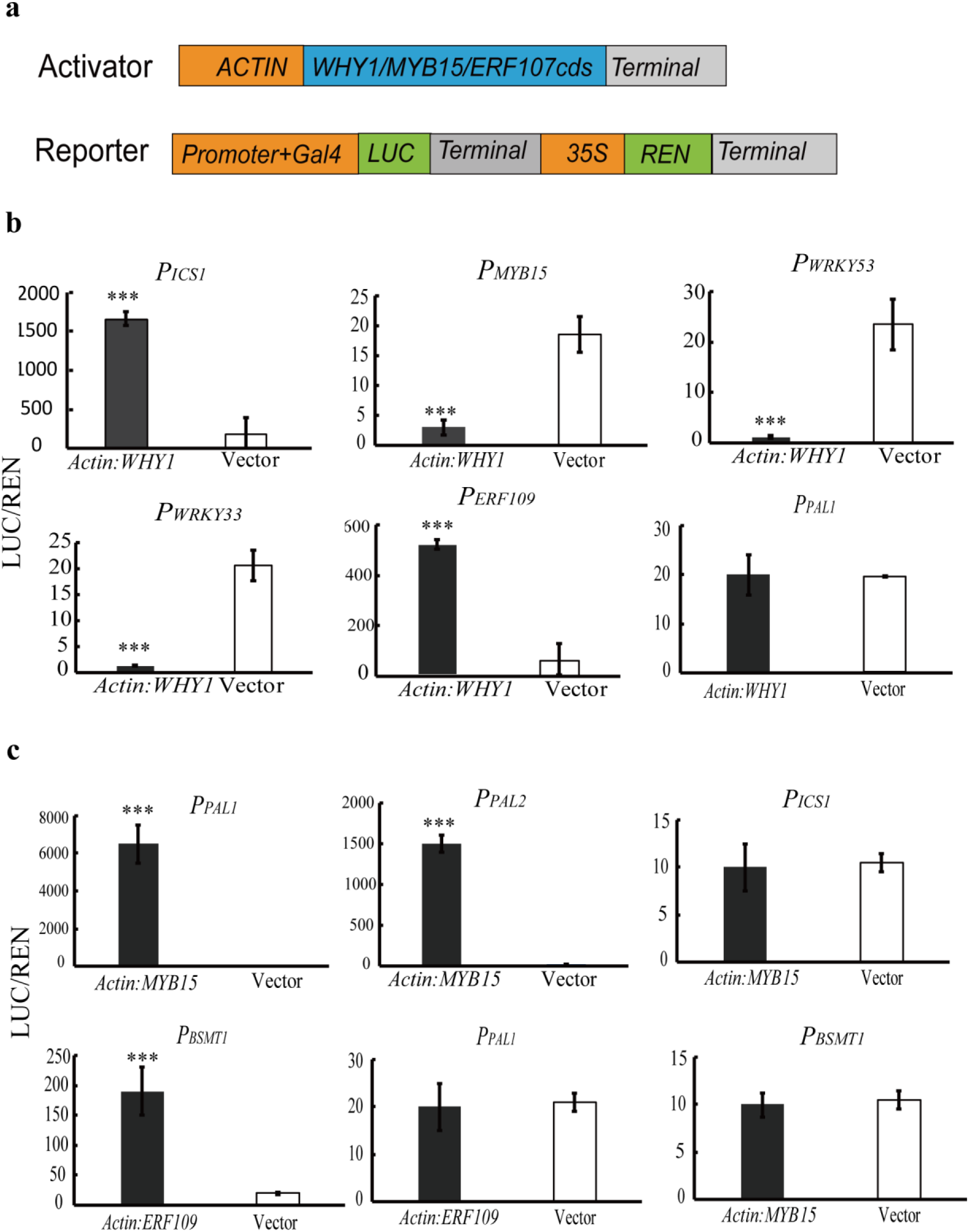
Promoter activation assays using the LUC/REN system a. Structure of activator and reporter constructs. b. The promoters of *ICS1, MYB15, ERF109, WRKY53*, and *WRKY33* genes are co-infiltrated with a vector containing WHY1 under the regulation of the ACTIN promoter. c, Co-infiltration of MYB15 and ERF109 with the *PAL1, PAL2, ICS1*, and *BSMT1* promoters. Background promoter activity is assayed by co-infiltration with an empty vector of the same type. Shown are means and SE of six biological replicates. Asterisks denote statistically significant differences from the empty vector calculated using Student’s t test: *, P, 0.05; **, P, 0.01; and ***, P, 0.001.

### WHY1 and MYB15/ERF109 regulate leaf senescence and ROS accumulation

Since WHY1 is a repressor of plant senescence at early stage (35-42 dag) of plant development (Miao et al. 2013), we compared the phenotype of the *pal1, sid2, myb15, erf109* mutants (Supplementary Fig S2) with the *why1* mutant to analyze if WHY1 effects on salicylic metabolism impact senescence. The phenotypes of the *pal1* and *sid2* plants have already been reported to delay senescence, and on the contrary, *oePAL1, oeSID2* and *bsmt1* plants showed an early senescence phenotype (Love et al., 2008; Rivas-San et al., 2011; Vlot et al., 2009; Huang et al. 2010). We analyzed all mutants with respect to a visible senescent yellow leaf ratio (Miao and Zentgraf, 2007) and reactive oxygen species (ROS) production by nitro blue tetrazolium chloride (NBT) staining assay and diaminobenzidine (DAB) staining assay under normal growth condition. The results showed that all of *pal1, sid2, myb15*, and *erf109* lines displayed a visible delayed senescence and less ROS production except for the *bsmt1* plants, which showed slightly earlier senescence and higher ROS accumulation similar to the *why1* and the *pWHY1* lines (Figure 6a-b).

**Figure 6.**
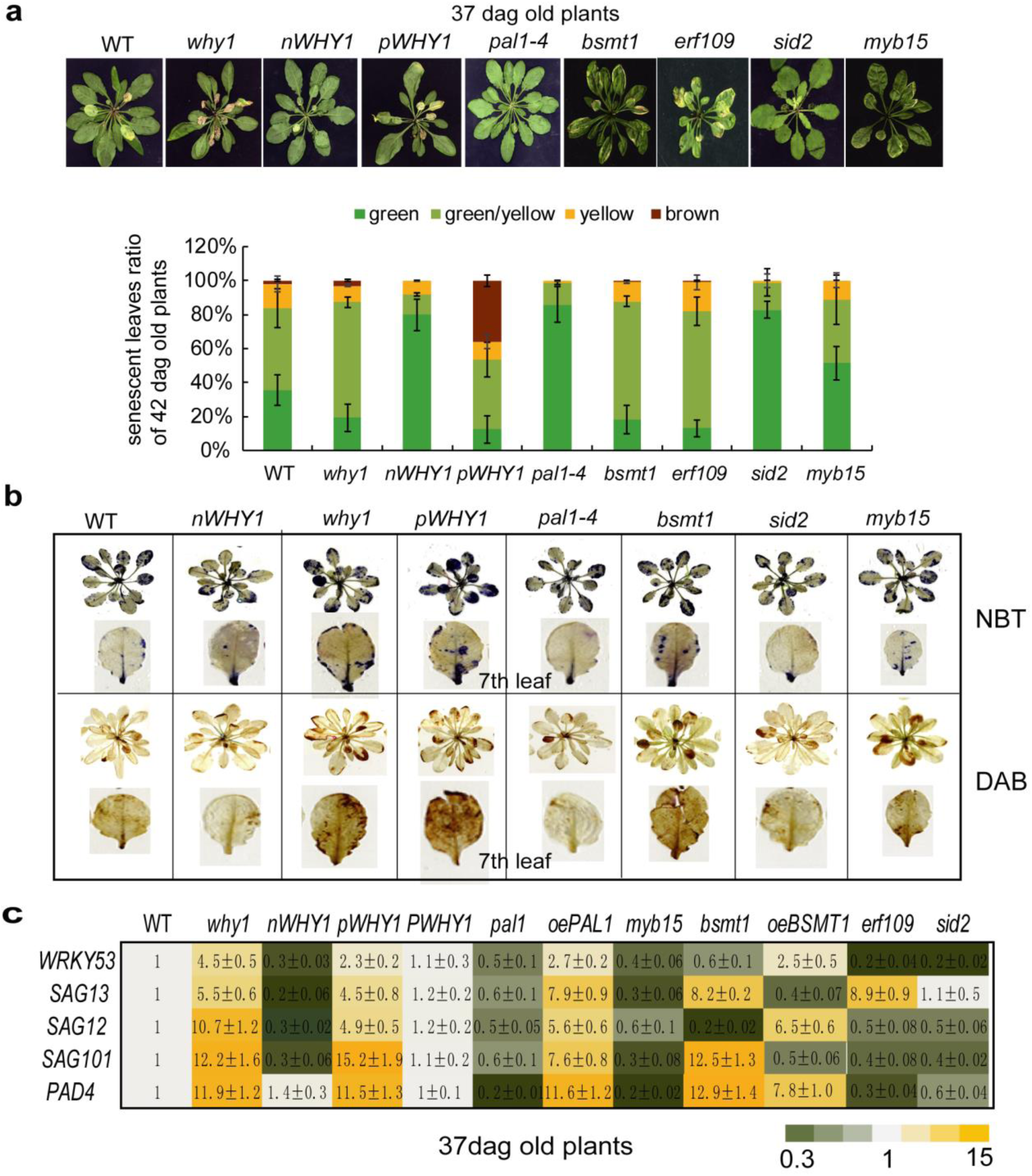
Phenotyping of loss- of *WHY1* and its downstream target genes mutants a. Phenotypes of loss-of *PAL1, ICS1, MYB15* and *BSMT1* at 37dag compared to *WHY1* mutants. Whole rosette (a-up) and senescent leaf ratio of 5 plants (a-down); b. ROS accumulation of loss-of *PAL1, ICS1, MYB15* and *BSMT1* at 37dag compared to *WHY1* mutants by NBT and DAB staining; c. The transcript levels of SAGs genes in the loss- or gain- of *PAL1, BSMT1* and loss- of *MBY15, ERF109* and *ICS1* plants at 37dag by qRT-PCR. The standard error is calculated from three biological replicates, the values are shown as means±SE. The wild-type at 37 dag was setup to 1 in the heatmap.

Furthermore, the transcript levels of senescence related genes *such as WRKY53, SAG12, SAG13, SAG101*, and *PAD4* were measured by qRT-PCR and indicated as heatmap (Figure 6c). They were upregulated in the *why1* and *pWHY1* plants,similar to the overexpressing *PAL1* (*oePAL1)* plants, however downregulated in the *pal1, myb15, and sid2* similar to the *nWHY1* plants (Figure 6c). Interestingly, in the overexpressing *BSMT1 (oeBSMT1)* line the transcript level of senescence related genes *SAG12* and *WRKY53* were upregulated, while the transcript level of *SAG13* and *SAG101* were downregulated, a reversed expression trend as compared to the *bsmt1* and *erf109* mutants (Figure 6c). However, the transcript level of *PAD4* was upregulated in the both *bsmt1* and *oeBSMT1*. This indicates that BSMT1 is involved in alternative signaling pathways between developmental senescence or stress related senescence

### SA level feedback affects the distribution of the WHY1 protein in plastids and the nucleus

WHY1 is required for SA- and pathogen-induced *PR1* expression (Desveaux et al. <u>2005</u>); WHY1 distribution is affected by protein modification (Ren et al. 2017) and cellular H_2_O_2_ level (Lin et al. 2019). To determine whether SA feedback would affect *WHY1* expression we quantified WHY1 transcription by qRT-PCR in response to exogenous MeSA in WT plants for 1, 4, 6, and 8 hours. Unexpectedly, MeSA treatment did not change the gene expression level of *WHY1* (Figure 7a). Thus, MeSA treatment probably affects WHY1 protein function or distribution in plastids or the nucleus. Thus, nuclear and plastid proteins isolated from 5-week-old WT rosettes after MeSA treatment for 4 hours were immunodetected with a specific monoclonal antibody against WHY1 (Lin et al. 2019; Supplementary Fig S4), and antibodies against Histone 3 and photosystem II (PSII) protein were used as markers for pure nuclear and plastid preparations (Figure 7b-c; Supplementary Fig S5). A water treatment served as control for MeSA application. Interestingly, the results now indicated that upon MeSA treatment for 4h, WHY1 accumulation significantly decreases in plastids and the nuclear isoform of WHY1 was altered in its status with small nWHY1 (29 kDa) levels slightly increasing, while large nWHY1 (37 kDa) levels were decreasing after MeSA treatment (Figure 7b-c). Thus, exogenous MeSA treatment affects WHY1 accumulation in plastids and alters the modification status of nWHY1 in the nucleus, a similar response as observed in response to H_2_O_2_ treatment (Lin et al. 2019). Furthermore, we analyzed WHY1 distribution between plastid and nucleus under the condition of SA deficiency. The nuclear and plastid fractions isolated from the single *sid2, pal1* mutants and double *sid2 pal1* mutant were subjected to immunoblotting using the WHY1 specific peptide antibody and the results demonstrate that pWHY1 in the *sid2, pal1* and *sid2 pal1* mutants significantly accumulated in plastids when compared to WT. Accordingly, the large nuclear WHY1 isoforms (37 kDa) were highly accumulating and the small nuclear WHY1 proteins (27 kDa) were declining in the *sid2* and *sid2 pal1* mutants, but not in the *pal1* single mutant (Figure b-d). This indicates that the ICS1 pathway plays a prominent role in modification of nWHY1 protein.

**Figure 7.**
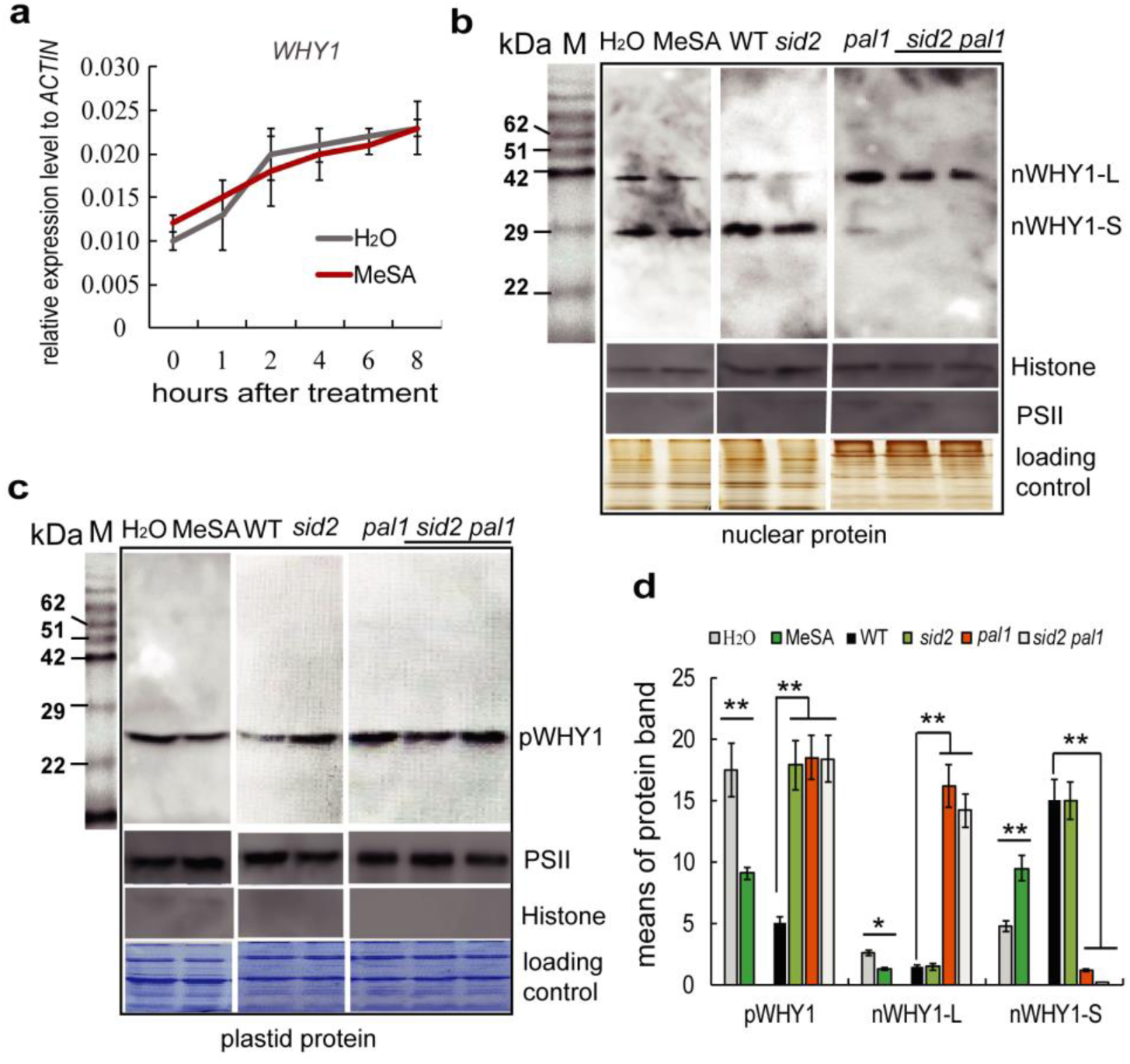
The plastid and nuclear isoform WHY1 protein immunodetection after the treatment of MeSA and in the *sid2, pal1* or double *sid2 pal1* mutants compared to WT a. The expression level of WHY1 in the WT plants after MeSA treatment for 1, 2, 4, 6, 8 hrs; b. WHY1 immunodetection in nuclear extracts after the treatment of MeSA for 4 hours, and in the *sid2, pal1* or double *sid2 pal1* mutants compared to WT; c. WHY1 immunodetection in plastid extracts after the treatment of MeSA for 4 hours, and in the *sid2, pal1* or double *sid2 pal1* mutants compared to WT. Coomassie and silver staining as the protein amount loading controls. L-WHY1: large size (37 kDa) of WHY1; S-WHY1: small size (29 kDa) of WHY1. The antibody against peptide WHY1 was prepared by company; d. The alteration of pWHY1 and nWHY1 after MeSA treatment or in the sid2, pal1 or double sid2 pal1 mutants compared to WT. The protein band signal is captured and calculated by Image J software program (http://www.di.uq.edu.au/sparqimagejblots). The data shows the average of three replicates. Asterisks (*P < 0.05, **P < 0.01) show significant differences to H_2_O treatment or WT according to Student’s t test.

## Discussion

It has become increasingly clear that dual location of proteins mediates diverse intercellular signaling processes, e.g. described for MAP kinase (Bobik et al. 2015; Chan et al. 2016), CIPK14 (Ren et al. 2017), but also hormone (ABA, SA) (Koussevitzky et al. 2007; Caplan et al. 2015; Kacprzak et al. 2019), or ROS (hydrogen peroxidase and singlet oxygen) signaling (Lin et al. 2019; Duan et al. 2019, Lv et al. 2019). Proteins with dual subcellular localization can affect transcription and display various functions in intracellular signaling (Lin et al., 2019; Isemer et al., 2012; Sun et al., 2011; Nevarez et al., 2017; Pesaresi and Kim, 2019; Wu et al., 2019; Woodson et al., 2011/2013). This study revealed that dual-located WHY1 protein directly activates *ICS1* expression in the nucleus at the late stage of plant development, and indirectly controls PAL1 and BSTM1 expression *via* alteration of *MYB15* and *ERF109* transcription at the early stage, thereby influencing the SA homeostasis in the cells during plant development. A SA level feedback affects in turn WHY1 distribution with a shift into the nucleus and preferential accumulation of the smaller 29 kDa form. This loop of nWHY1 integrating SA homeostasis via PAL1/ICS1 and BSMT1 plays a pivotal role in controlling leaf senescence.

Elucidation of biosynthesis and catabolism of SA is important for understanding its biological functions. 10% of SA is synthesized either from L-phenylalanine via the PAL pathway in the cytoplasm or up to 90% from chorismate via ICS1/SID2 (ISOCHORISMATE SYNTHASE1/SALICYLIC ACID INDUCTION DEFICIENT2) in chloroplasts, the latter of which is responsible for the bulk of SA produced during pathogen infection in *Arabidopsis* (Dempsey et al. 2011). Endogenous SA can also undergo a series of chemical modifications including hydroxylation by salicylate hydroxylase (Yamamoto et al. 1965; Zhang et al., 2013), glycosylation by glycosyltransferases (Lim et al. 2002; Dean et al. 2008), methylation by BSMT1 (Park et al., 2007) and amino acid or sugar conjugation by XXX (Zhang et al., 2007; Bartsch et al. 2010). The microarray data and qRT-PCR results show that the gene expression levels of developmental related transcription factors were upregulated, and that of stress-related gene were downregulated in the *why1* plants (Figure 3; supplementary dataset 1-4). The expression levels of *ICS1, PAL1* and *BSMT1* were altered significantly in the *why1* mutant during plant aging (Figure 1); this alteration can be rescued completely by complementation of nWHY1 and pnWHY1 (Figure 2). As we knew, nWHY1 could directly bind to the promoters of many targeted genes such as *WRKY53, S40, Kenisin, PR10a* (Desveaux et al. 2005; Miao et al. 2013; Krupinska et al. 2017; Xiong et al. 2009), as well as *MYB15, MYC1/2, ICS1* and several ERF family members from our WHY1 ChIP-seq dataset (Figure 4; Miao et al., 2013). The nWHY1 represses most of downstream developmental related target gene expression such as *WRKY53, WRKY33, MYB15, TTG2 etc.* (Figure 3; Supplementary dataset). However, it can also promote expression of many stress-related genes such as *HvS40 (*Krupinska et al. 2013*), PR1* (Desveaux et al. 2005), redox responsive transcription factors (Foyer et al. 2014), *ICS1*, and *ERF109* (Figure 5; Figure 3; Supplementary dataset). Several MYB family members can bind to the promoter of *PAL1/PAL2* (Battal et al. 2019), and among these, MYB15 was shown to bind to the promoter of *PAL1* and *ICE1* promoter by ChIP-qPCR. MYB15 mainly plays a virtual role in immunity and cold response (Chezem et al. 2017; Kim et al. 2017; Wang et al. 2019). Our results further confirmed that MYB15 could activate *PAL1* expression. ERF-binding cis elements are enriched in the promoter region of *BSMT1*. However, *ERF109* as a target gene of WHY1, which was identified in our ChIP-seq dataset (Miao et al. 2013; Figure 4), was reported not to bind to the promoter region of *BSMT1* as shown in yeast one hybrid and gel shift assays (Ximiao Shi, Master thesis, 2018). In contrast, ERF109 can activate *BSMT1* expression in our LUC/REN transit assay (Figure 5), supporting our ChIP-seq data. The *erf109* and *bsmt1* mutants accumulate high levels of anthocyanin in response to high light (Foy et al. 2015), but the regulatory mechanism is currently unknown. Therefore, the balance module of nWHY1/MYB15-PAL1 and nWHY1/ERF109-BSMT1 at early stage (37 dag) and WHY1/ICS1 regulation at late stage (42 dag) determines SA homeostasis during plant development. The imbalance of PAL1/BSMT1 activity at 37 dag in the *why1* mutant and repression of ICS1 at 42 dag of plant development may result in earlier SA accumulation for about one week. Thus, nWHY1 impacts the SA homeostasis *via* mediating PAL1 or ICS1 and BSMT1 activity in the cells during plant aging.

The WHIRLY family is considered to associate with retrograde signaling. Due to their dual-location and function in the nucleus and plastids (Krause et al., 2009), it has been supposed that WHIRLY1 could move from plastid to the nucleus (Isemer et al., 2012). The plastid isoform of WHIRLY1 affects the *miRNA* biogenesis in the nucleus (Swida-Barteczka et al. 2018). Previously, we showed that the WHY1 protein can be phosphorylated by CIPK14 kinase or oxidized by H_2_O_2_, leading to different subcellular localization in the nucleus or in plastids, respectively (Ren et al. 2017; Lin et al. 2019). Here, we show that loss- of WHY1 results in five days earlier SA production during plant development, thereby accelerating plant senescence. Complementation with pWHY1, did not revert the SA accumulation phenotype. On the contrary, the pWHY1 further increased SA accumulation during plant development. Consistently, gene expression of *PAL1* is promoted, while that of *BSMT1* is repressed at 37 dag, while ICS1 is activated at 42 dag (Figure 1). This phenomenon can be explained by two mechanisms: 1) H_2_O_2_ is known to affect SA levels via the ICS1 pathway (Leon et al. 1995; Dat et al. 1998; Chaouch et al. 2010; Guo et al. 2017) and recent data link pWHY1 to ROS production *via* photosystem I/II (PSI/PSII) (Huang et al. 2017; Lin et al. 2019). Thus, pWHY1 might increase SA level at 42 dag by modulation of the ICS1 pathway via photosystem induced ROS accumulation to cause an early senescent phenotype. 2) pWHY1 coordinating SA homeostasis is feedback controlled by cellular SA levels ((Desveaux et al. 2005; Isemer et al. 2012; Caplan et al. 2017), so this WHY1 isoform changes from plastid to nucleus repressing *MYB15* and *PAL1* expression (Huang et al. 2010; Duan et al., 2019) and activating *ERF109* and *BSMT1* expression in response to stress cues such as high light (Estavillo et al. 2011). This demonstrates that dual located pWHY1/nWHY1 affects SA homeostasis most likely *via* connection with PSI/II mediated ROS affecting leaf senescence.

The distribution of WHY1 between plastids and the nucleus depends not only on its modification status (Ren et al. 2017) but also on environmental cues or cellular signals such as H_2_O_2_ (Lin et al. 2019) and SA (this work). Though, the SA signal cannot promote CIPK14 expression (unpublished data), MeSA treatment feedback alters the nWHY1 protein status (37 kDa or 29 kDa form) (Figure 7) similar to barley WHY1 (Grabowski et al. 2008) and nWHY1 after treatment with H_2_O_2_ in *Arabidopsis* (Lin et al. 2019). The nature of modification resulting in both forms is yet unknown and has to be revealed in future. More interestingly, MeSA treatment reduced WHY1 accumulation in plastids, which stands in contrast to H_2_O_2_ treatment (Lin et al. 2019). These phenomena are further elucidated in the SA deficient mutants such as *pal1, sid2* and double *pal1 sid2* mutants. Loss-of ICS1 (*sid2*) decreases the modified state of nWHY1 level, while loss-of ICS1 or PAL1 increases WHY1 accumulation in plastids. It is known that ICS1 is located in plastids and is responsible for the bulk production of SA in response to salt or pathogens (Kumazaki and Suzuki, 2019). Plastid-derived SA can be transported from plastid to the nucleus *via* stromule (Caplan et al. 2015). It is speculated that this kind of SA might influence the nuclear isoform of WHY1, which small form (29 kDa) activates the stress related gene expression, such as *S40, ICS1* (Krupinska et al., 2013; Figure 4-5), while the large form (37 kDa) represses gene expression, as shown for *WRKY53, WRKY33, MYB15* (Miao et al. 2013; Figure 4-5). Furthermore, it has been reported that phosphorylation of WHY1 by CIPK14 promoted its binding affinity at the promoter of *WRKY53* and *WRKY33* and repressed *WRKY53* and *WRKY33* expression (Ren et al. 2017) and that CIPK kinase expression level rapidly increased in response to salt or pathogen, accompanying increasing Ca^2+^, H_2_O_2_ and SA levels in the cells (Sardar et al., 2017).

## Conclusion

We conclude that WHY1 exerts dual functions in plastids and the nucleus. Nuclear WHY1 maintains SA homeostasis by directly affecting *ICS1* and indirectly affecting *PAL1* and *BSTM1* expression via *MYB15* and *ERF109*. The pWHY1 isoform promotes PAL1/ICS1 expression and represses BSMT1 facilitating high SA accumulation, resulting in early senescence, similar to *bsmt1* mutants. Interestingly, MeSA treatment altered the nWHY1 status (increasing the 29 kDa form of WHY1, while decreasing the 37 kDa form), going along with declined pWHY1 accumulation. These results indicate that pWHY1/nWHY1 distribution in the nucleus and chloroplast allows balancing SA and H_2_O_2_ homeostasis, in a developmental dependent manner, thereby affecting leaf senescence in *Arabidopsis* (Figure 8).

**Figure 8.**
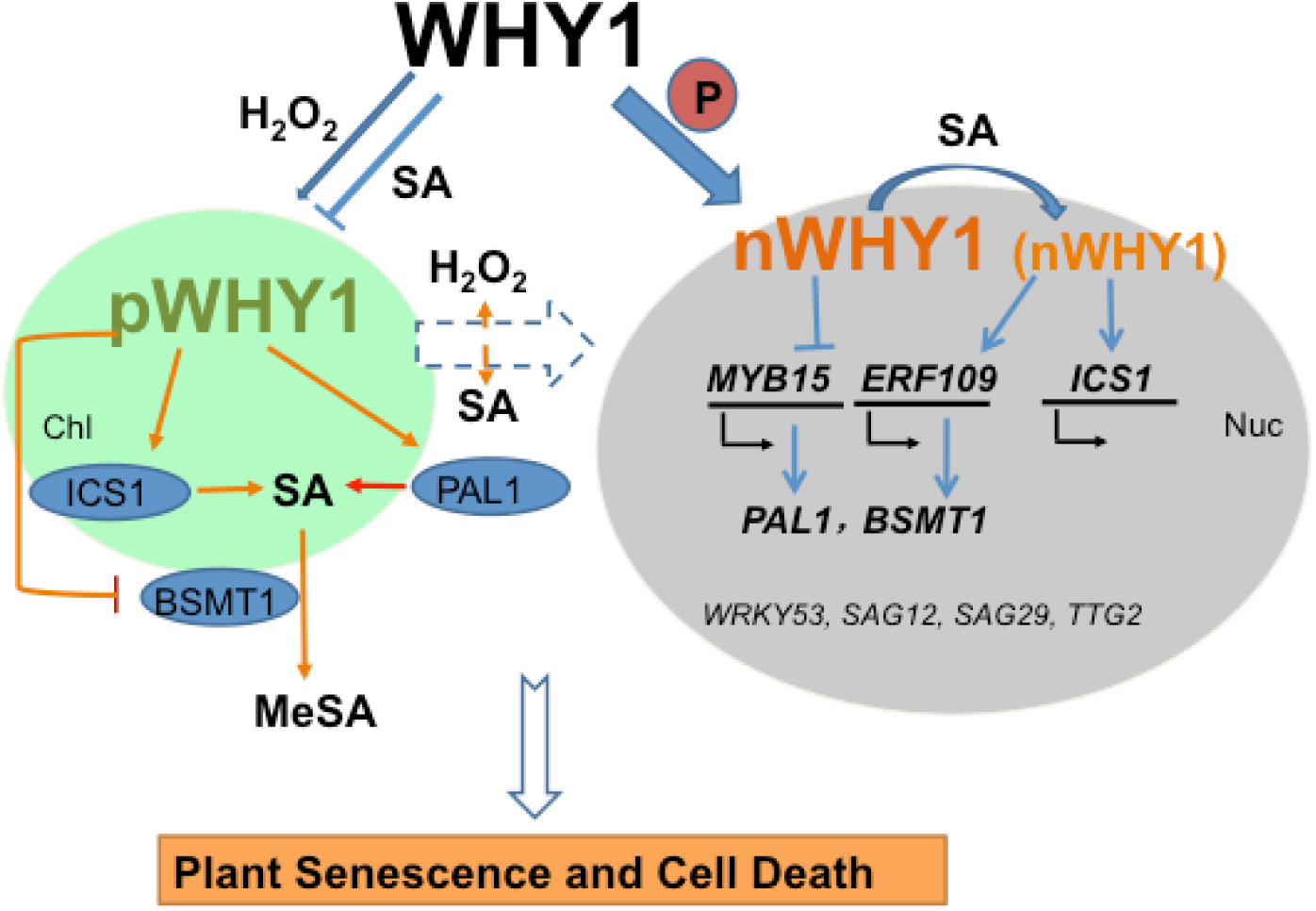
A working model of the senescence pathway performed by the dual located WHY1 in response to SA. The nuclear isoforms of WHY1 are represented as both a large molecular mass (37 kDa, bigger letters in the Figure) and a small molecular mass (29 kDa, smaller letters). WHY1 has dual functions in plastids and the nucleus. Loss of WHY1 increases SA accumulation at early stage (37 dag) through increasing PAL1 expression and repressing BSMT1; Elevated SA promotes nuclear WHY1 de-modification and promotes ICS1 and BSMT1 expression thereby balancing SA homeostasis in the cells. High SA levels by ICS1 cause feedback enhancing ROS accumulation, promoting senescence. pWHY1 stimulates PAL1/ICS1 expression but represses BSMT1, allowing high levels of SA, leading also to early senescence. Thus, distribution of WHY1 organelle isoforms and the putative feedback of SA form a circularly integrated regulatory network during plant senescence in a developmental dependent manner. Plastid (Chl) is shown as a green ovary, nucleus (Nuc) as a grey ovary, lines for regulation, fat arrows for transfer or translocation, broken lines for uncertainty.

## Materials and Methods

### Plant materials

All *Arabidopsis thaliana* mutants are in Col-0 background. The T-DNA insertion lines *why1* (Salk_023713), *sid2, pal1, bsmt1* (SAIL_776_B10), *myb15* (*myb15-1* SALK_151976, *myb15-2* SK2722) were kindly provided by other scientists; The *erf109* (SALK_150614) and over-expression lines of ERF109 gene (CS2102255) were obtained from the Nottingham Arabidopsis stock center (NASC). Homozygous plants were selected and confirmed by PCR or RT-PCR using gDNA and mRNA as templates (Supplementary Fig S2), respectively (http://signal.salk.edu/tdnaprimers.2.html). The overexpressing *nWHY1-HA* lines that produce the WHY1 protein located only in the nucleus, the overexpressing *pnWHY1-HA* lined that produce the WHY1 protein dually located in plastids and the nucleus, the complement *PWHY1-HA* (*Pwhy1:pnWHY1-HA*) line, and the *pWHY1-HA* lines that harbor the construct of the full length WHY1 plus nuclear export peptide sequence fused to HA-tag produces WHY1 protein located only in plastids have been constructed in our lab (Miao et al. 2013; Lin et al. 2019).

Seeds are germinated on wet filter paper followed by vernalization at 4°C for 2 d, then transplanted to vermiculite and are grown in a climatic chamber (100 μE/h, 13h of light at 22°C/11h of dark at 18°C, 60% relative humidity). The rosette leaves are labeled with colored threads after emergence, as described previously (Hinderhofer and Zentgraf 2001).

For MeSA treatment, rosette leaves are collected at 1, 2, 4, 6, and 8 hours after spraying with 100 µM MeSA and stored in liquid nitrogen or −80 °C for later use in RNA or protein isolations. Mock treatments used distilled water instead.

### Measurement SA contents in rosette leaves

SA was extracted from 0.2 g the 5^th^ leaf from individual plants at different stages of development and measured by reversed-phase high-performance liquid chromatography (HPLC) on an Agilent1260 system with a C18 column as previously described (Verberne et al. 2002) with small modifications: SA was thoroughly separated from the complex mixture by methanol containing 10% of sodium acetate with pH 6.0 (Lin et al. 2017). Fluorescence detection (excitation at 305 nm and emission at 407 nm) was applied and 3-Hydroxybenzoic acid (3-HBA) was used as an internal standard (Aboul-Soud et al. 2004). Conjugated and free SA was detected at the same time. Three independent biological replicates were performed for each data point.

### Staining of ROS

Visualization of H_2_O_2_ accumulation in leaves was performed using the 3’,3’-diaininobenzidine (DAB) staining method according to Zhang et al. (2014) and Huang et al. (2019). Detached rosette leaves were vacuum filtered in 20 mL staining solution containing 1 mg/mL DAB in 50 mM Tris-HCl, pH 5.0 for 10 min, and incubated in the darkness at room temperature for 12 h. The leaves were destained by boiling in a mixture of ethanol, glycerol and acetic acid (3/1/1, v/v/v) for 15 min before imaging.

Detection of superoxide free radicals were performed by the nitroblue tetrazolium (NBT) staining method as described in Lee et al. (2002). The whole rosette leaves of 5- to 6-week-old plants were harvested and immersed in 0.1 mg ml-1 NBT solution (25 mM HEPES, pH7.6). After vacuum infiltration, samples were incubated at 25°C for 2 h in the darkness. Subsequently stained samples were bleached in 70% ethanol and incubated further for 24 h at 25°C to remove the chlorophyll.

Imaging was conducted using an Epson Perfection V600 Photo scanner (Epson China, Beijing, China).

### Quantitative real-time PCR analysis (qRT-PCR)

The qRT-PCR was performed using SYBR Green master mix (SABiosciences, Frederick, MD, USA) according to the manufacturer’s instructions. Complementary DNA synthesis was carried out using a Fermentas first-strand complementary DNA synthesis kit (Thermo Fisher Scientific, Waltham, MA, USA) on RNA from 28-55-day-old plants grown under normal light conditions. Complementary DNAs were diluted 20-fold prior to quantitative PCR experiments. The Touch 1000 platform (Bio-Rad) was used for qRT-PCR experiments, and the data were analyzed using Bio-Rad software version 1.5. We used *GAPC2* or *ACTIN* as internal reference genes for calculation of relative expression. Primers are listed in Supplemental Table S1. All determinations were conducted in three biological replicates.

### Isolation and detection of plastid and nuclear proteins

Chloroplasts and nuclei were prepared and purified as described previously (Ren et al. 2017). Approximately 10 microgram proteins of each fraction was separated on 14% (w/v) polyacrylamide gels. After transfer to nitrocellulose membranes, immunodetection followed using specific antibodies against the WHY1 C-terminal peptide CASPNYGGDYEWNR (Faan, Hangzhou, China). To monitor the purity of the chloroplast and nuclear fractions, we used antibodies against the cytochrome b559 apoprotein A or the histone H3 (Cell Signaling, Munich, Germany), respectively (Lin et al. 2019).

### ChIP-qPCR assay

Four-week-old rosettes of transgenic plants expressing the *Pwhy1:WHY1-HA* to complement the *why1* knockout background were used for sample preparations. The cross-linked DNA fragments ranging from 200 to 1000 bp in length were immunoprecipitated by an antibody against the HA-tag (Cell Signaling, Munich, Germany). The enrichments of the selected promoter regions of both genes were resolved by comparing the amounts in the precipitated and non-precipitated (input) DNA samples, which were quantified by quantitative PCR using designed region-specific primers (Supplementary Table S1 and Figure 4). Material from the *why1* mutant served as a mock control and was used for normalizations to calculate the fold enrichment. The experiments were performed three times biological replicates.

### Cloning and Construction of Vectors

The promoter sequences of 2kb upstream of ATG of *MYB15* and the *ERF109, WRKY53, PAL1, PAL2, ICS1*, and *WRKY33* genomic sequence were PCR-amplified and then restricted with *Kpn*I and *Xho*I, or *Xho*I and *Pst*I respectively and sub-cloned into the pFLAP vector The entire cassette was then excised with *Kpn*I and *Asc*I and cloned into the binary vector pBIN +.

For dual Luciferase assays, promoter sequences were PCR-amplified, digested with *Nco*I and *Kpn*I and cloned into the pGreenII 0800-LUC binary vector (provided by Roger P. Hellens). DNA constructs used for *N. benthamiana* agro-infiltration and for agrobacteria-mediated plant transformation were constructed with the Goldenbraid cloning (Sarrion Perdigones et al., 2013).

*MYB15, ERF109* and *WHY1* coding sequences were subcloned into a pUPD vector. In the dual Luciferase assays; MYB15, ERF109, and WHY1 were in the 1α1 vectors which are based on a pGREENII backbone. For generating the genes overexpression construct, a CDS fragment was amplified subcloned into pGEM-T Easy (Promega), excised with *Bam*HI and *Sal*I restriction enzymes and cloned under the CaMV-35S promoter into pFLAP, before restriction with *Pac*I and *AscI* and ligation to the pBIN+ binary vector.

### Dual-luciferase activity assay

*Nicotiana benthamiana* plants were grown in climate rooms (22°C, 16/8 h of light/dark). Plants were grown until they had six leaves and then infiltrated with *Agrobacterium tumefaciens GV3101*. Plants were maintained in the climate rooms and, after 4 to 5 d, 1-cm discs were collected from the fourth and fifth leaves of each plant. Six biological replicates with their respective negative controls were used per assay. The experiment was performed as previously described (Hellens et al., 2005) with minor changes. Agrobacterium was grown over night in LB and brought to a final O.D.600 0.2 in infiltration buffer. Co-infiltrated Agrobacterium carried separate plasmids; 900 μl of an empty cassette or one that contains the transcription factor driven by the tomato 2 kb UBQ10 (SOLYC7G064130) promoter region, and 100 μl of the reporter cassette carrying one of the test promoters. Leaf discs were homogenized in 300 μl of a passive lysis buffer. 25 μl of a 1/100 dilution of the crude extract was assayed in 125 μl of Luciferase assay buffer, and LUC and REN chemiluminescence of each sample was measured in separate wells on the same plate. RLU were measured in a Turner 20/20 luminometer, with a 5 seconds delay and 15 seconds measurement. Raw data was collected and the LUC/REN ratio was calculated for each sample. Biological samples were polled together and a student’s t-test was performed against a background control for each experiment as described in the results section. The entire experiment was repeated a second time under similar conditions to confirm the regulatory effect of transcription factors.

### Microarray Analysis

Two biological replicates were sampled from leaves of wild-type, *VEX:pWHY1/why1, VEX:nWHY1/why1*, and the *why1* plants (see our previous paper Ren et al. 2017). Extracted RNA was then amplified and labeled using the standard Affymetrix protocol and hybridized to Affymetrix ATH1 GeneChips according to the manufacturer’s guidelines (Katari et al. 2010). Statistical analysis of transcriptome data was carried out using Parke Genome Suite software (www.partek.com). Data preprocessing and normalization were performed using the Robust Microarray Averaging algorithm (Irizarry et al., 2003). Batch effects between the replicates were not found. Differentially expressed genes were identified by using ANOVA according to false discovery rate, p-value 0.05 and at least a 2-fold change between the genotypes (Supplementary Dataset 1-4).

### Statistical analysis

Quantitative data were determined by at least three biological replicates and the statistical significance was analyzed either using two-way ANOVA or pair-wide multiple t-tests, with the GraphPad Prism software (version 7).

## Acknowledgments

We acknowledge Dr. Hongwei Guo (University of Southern Technology) for providing the *sid2*, Dr. Zhixiang Chen (Purdue University) for providing the *pal1-4* seeds, Dr. Nicola Clay (Yale University) for providing the seed of *oeMYB15* and the *myb15*, Dr. Daniel Klessig (University of California, Berkeley) for providing the *bsmt1* seeds. We acknowledge the European Arabidopsis Stock Centre providing series of Arabidopsis mutant seeds (*oe:ERF109* and *erf109* lines). This work is supported by a grant from the Natural Science Foundation of China (NSFC, No. 31770318, 31470383), and by a grant from Fujian Provincial NSF (No. 2016J01103), and by a grant of international exchange program of Fujian Agriculture and Forestry University (KXB16009A).

## Author contributions

Y.M. designed the study. W.F.L. performed SA measurements, western blots, phenotyping, and qRT-PCR. D.H. performed ChIP-seq, ChIP-qPCR. H.Z. performed plasmid constructs and promoter activation activity and the mutants screening. B.H.W performed microarray data analyses. W.F.L. and Y.M. analyzed the data. Y.M. and D.S. wrote the paper. D.C. critically read the paper.

## Competing interests

The authors declare no competing interests.

## Supporting information

Supplementary Fig S1 Transcript levels of *ICS2, UGT71B1, UGT89B1, UGT74F2* and *S3H* in the *why1* mutant compared to WT during plant aging

Supplementary Fig S2 Verification of mutant plants used in this study..

Supplementary Fig S3 Venn analysis of transcriptome of nWHY1, pWHY1, pnWHY1 and *why1*

Supplementary Fig S4 Antibody specificity test.

Supplementary Fig S5 Western blot detection to certify purity of nuclear protein and plastid protein extracts.

Supplementary Table S1. The list of primer sequences for PCR in this study Supplementary dataset 1-4

